# Bayesian Inference of the Evolution of a Phenotype Distribution on a Phylogenetic Tree

**DOI:** 10.1101/040980

**Authors:** M. Azim Ansari, Xavier Didelot

## Abstract

The distribution of a phenotype on a phylogenetic tree is often a quantity of interest. Many phenotypes have imperfect heritability, so that a measurement of the phenotype for an individual can be thought of as a single realisation from the phenotype distribution of that individual. If all individuals in a phylogeny had the same phenotype distribution, measured phenotypes would be randomly distributed on the tree leaves. This is however often not the case, implying that the phenotype distribution evolves over time. Here we propose a new model based on this principle of evolving phenotype distribution on the branches of a phylogeny, which is different from ancestral state reconstruction where the phenotype itself is assumed to evolve. We develop an efficient Bayesian inference method to estimate the parameters of our model and to test the evidence for changes in the phenotype distribution. We use multiple simulated datasets to show that our algorithm has good sensitivity and specificity properties. Since our method identifies branches on the tree on which the phenotype distribution has changed, it is able to break down a tree into components for which this distribution is unique and constant. We present two applications of our method, one investigating the association between HIV genetic variation and human leukocyte antigen, and the other studying host range distribution in a lineage of *Salmonella enterica*, and we discuss many other potential applications. All the methods described in this paper are implemented in a software package called TreeBreaker which is freely available for download at https://github.com/ansariazim/TreeBreaker

## Introduction

Understanding phenotypic variations and their relative association with genotypic variations is one of the central aims of molecular biology. The expression of a phenotype is usually dependent on both genetic and environmental factors, with heritability measuring their relative importance [1]. When the heritability is non-zero, genetically similar individuals are more likely to have similar phenotypes, and this is especially relevant for species that reproduce clonally, so that closely related individuals are virtually identical genetically. However, genotype-phenotype maps are usually complex and phenotypic plasticity means that phenotype expression can differ even for genetically identical individuals due to dependency on environmental factors [2, 3]. Conversely, observing closely related individuals with the same phenotype does not necessarily imply a low importance of environmental factors, since close relatives are also likely to live in the same environmental conditions [1]. The same effect also occurs in sexually reproducing species as evolutionary forces such as spatial population structure, environmental pressures and inbreeding result in groups within which individuals are more genetically homologous, and therefore more phenotypically similar, than individuals from different groups [4, 5].

To understand the relationship between a phenotype and a genotype, it is necessary to investigate how the phenotype is distributed according to genotypic values. This requires to quantify how the genotypes are related to each other which is often achieved using phylogenetic trees [6]. For clonal organisms, the tree may represent the clonal genealogy of how individuals are related with one another for non-recombinant regions [7, 8]. For sexual organisms, the phylogenies may be built for individual genomic loci, resulting in so-called gene trees by contrast with the species tree which contains them [9]. Visual inspection of a phylogenetic tree with tips annotated by phenotypes gives a first impression of their relationship, and this type of figure features heavily in the molecular biology literature of both clonal and sexual organisms. A more quantitative approach is however needed if the tree is too large to be shown, the interesting patterns too subtle to be seen, or to estimate evolutionary parameters and test competing hypotheses.

Phylogenetic comparative methods can be used, for example to test the phylogenetic signal in a phenotype [10, 11] or to compare the association between two phenotypes given the phylogeny [12], but do not provide a complete description of the phenotype distribution on a tree. Ancestral state reconstruction of the phenotype given the tree [13, 14] is often used for this and can provide quantitative insights, for example an estimate of the phenotypic evolutionary rate. The maximum likelihood approach to ancestral state reconstruction [15] has been extended in many ways by refining the model of phenotypic evolution on the tree, for example allowing to detect branches where the phenotypic evolutionary rate changes [16, 17]. However, ancestral state reconstruction is problematic for any phenotype with imperfect heritability: identical genotypes can then have different phenotypic values, implying an infinitely high rate of phenotypic evolution between them which is not biologically meaningful. Other difficulties arise if the phylogeny is imperfectly reconstructed or the phenotype inaccurately measured, which is always a possibility. Consequently, ancestral state reconstruction does not always provide reliable results, for example when applied to phylogeography [18].

When heritability is not complete, a phenotypic measurement can be seen as just one realisation from the phenotypic distribution of a given individual, with this distribution being what evolves on the tree rather than the phenotypic measurement itself. Based on this idea, here we present a novel Bayesian statistical method which takes as input a phylogenetic tree and discrete tip phenotype measurements, and identifies the branches on which the phenotype distribution has changed. The tree is therefore divided into monophyletic and paraphyletic groups that have unique distributions over the phenotype space. We also perform Bayesian hypothesis testing [19] to assess whether there is evidence for different parts of the tree having distinct phenotype distributions. We build a stochastic model in which changepoints occur on a phylogenetic tree [20], each of which affects the distribution of observed phenotype for the descendent leaves. Careful parametrisation enables the use of a fixed-dimension Monte-Carlo Markov Chain (MCMC) algorithm [21] to sample from the posterior distribution of the model parameters, and we reserve reversible jumps [22] to compare the model with a model without any changepoint. In the following sections we present our model, inference procedure and the results of simulation studies to measure the sensitivity and specificity of our method. Finally we present the application of our method to two real datasets in HIV evasion and bacterial ecology.

## Model and Methods

### Description of the model

We consider that changepoints happen as a Poisson process with rate λ on the branches of the input tree. For a phenotype with *K* categories, we model each changepoint event as a new probability mass function ***q*** = (*q*_1_,…,*q*_*K*_) which specifies the probability of having each of the *K* phenotypes for the individuals affected by the changepoint. Figure 1 illustrates the model for *K =* 2. The observed phenotype of each individual is shown on the tips of the tree which are coloured as black and red. Changepoints have happened on three branches which divided the tree into four sections (white, blue, green and yellow). All individuals in the same section have the same distribution ***q*** over the phenotype space.

**Figure 1.**
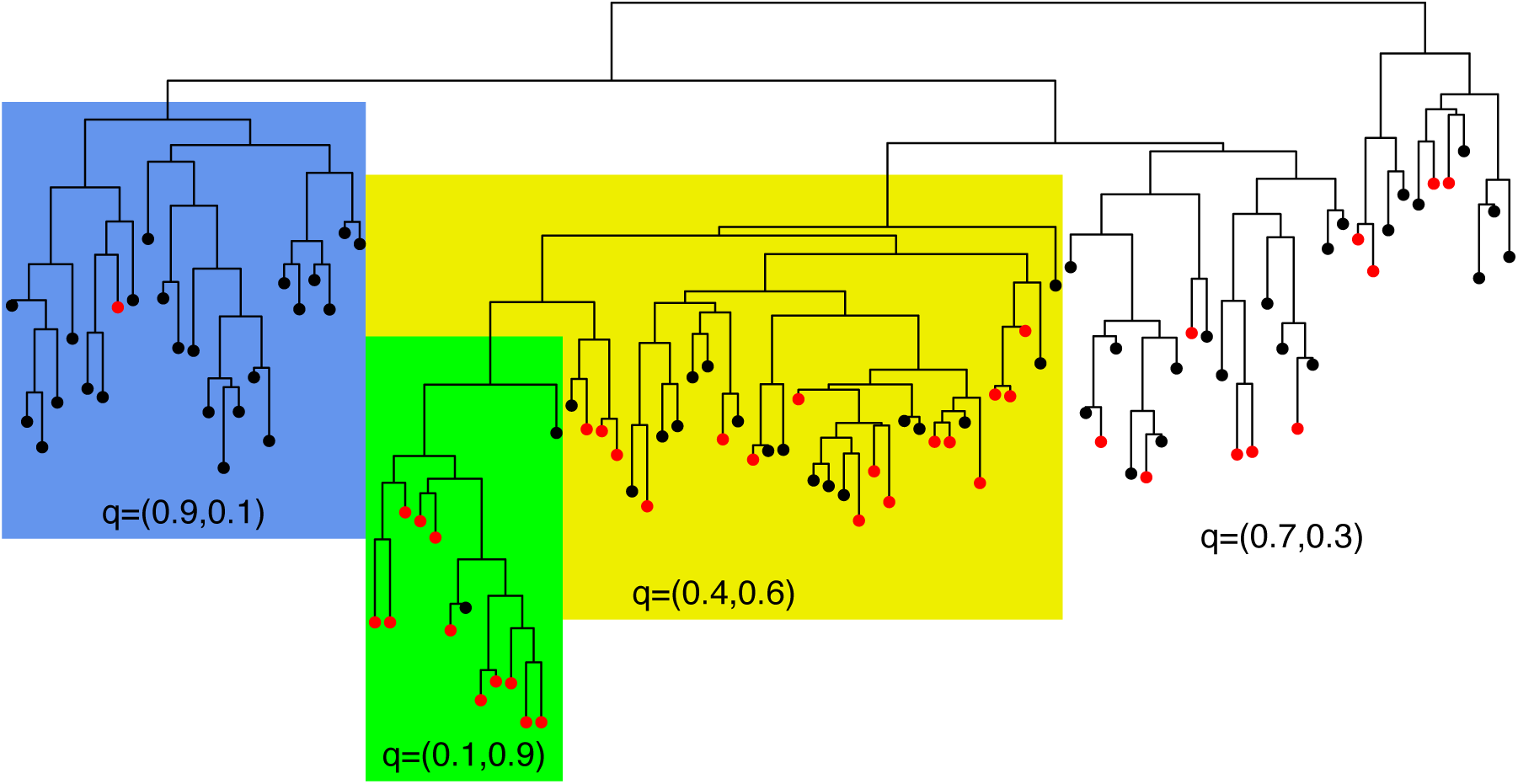
Illustration of the model. Changepoints occurred on three branches, which divided the tree into four sections (white, blue, green and yellow), each of which has different probabilities of the first (black) and second (red) phenotypes.

Let *N* and *B* denote the number of tips and branches in the tree, respectively (if the tree is bifurcating then *B* = 2*N*−2). We define ***b*** = (*b*_1_, …, *b*_*B*_) as a binary vector with *B* elements which represent the branches of the tree. If branch *i* holds at least one changepoint, then *b*_*i*_= 1 else *b*_*i*_ = 0. Let m denote the number of sections of the tree divided according to ***b*** (Figure 1), the likelihood of the observed phenotypes of the individuals *D* is given by:

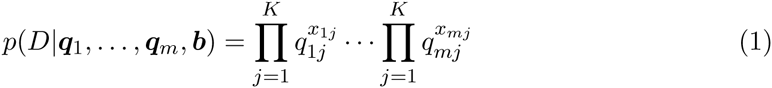

where ***q***_*i*_ = (*q*_*i*1_,…,*q*_*iK*_) and *q*_*ij*_ gives the probability that an individual in section *i* expresses phenotype *j*, so that 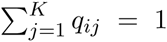 for *i* = 1, …, *m*. We also define *x*_*i*_ = (*x*_*i*1_,…,*x*_*iK*_) where *x*_*ij*_ is the number of observed individuals in section *i* which have expressed phenotype *j*, so that 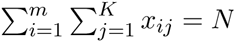.

The prior probabilities of branch *i* of length *l*_*i*_ having no or at least one changepoint are respectively equal to Pr(*b*_*i*_ = 0|λ) = *e^-λli^* and Pr(*b*_*i*_ = 1|λ) = 1 - *e^-λli^*, so that:

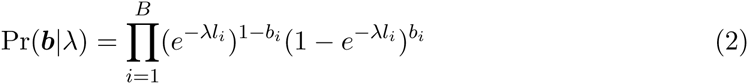

We consider a flat Dirichlet prior for all *q*_*i*_ such that *p*(***q***_*i*_) = Γ(*K)*, and an exponential prior on λ with parameter 1/T where 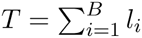 is the sum of the branch lengths of the tree. This implies a parsimonious prior expectation of one for the number of changepoints on the tree.

We are now in a position to describe the posterior distribution of the model parameters ***q***_*i*_,…, ***q***_*m*_, ***b*** and λ:

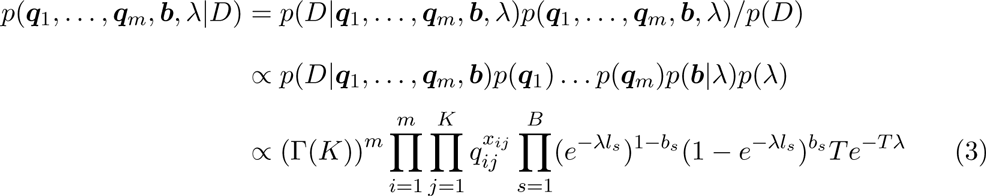

The dimensionality of the model parameters changes with ***b***. If ***b*** divides the tree into two sections then there are four parameters ***(q***_1_, ***q***_2_, ***b***, λ) in the model whereas if ***b*** divides the tree into three sections then there are five parameters (***q***_1_, ***q***_2_, ***q***_3_, ***b***, λ) in the model. This could potentially be addressed using reversible jumps [22]. Instead we marginalise all the ***q***_*i*_ which results in a fixed dimension model. The marginal posterior density for ***b*** and λ is given by:

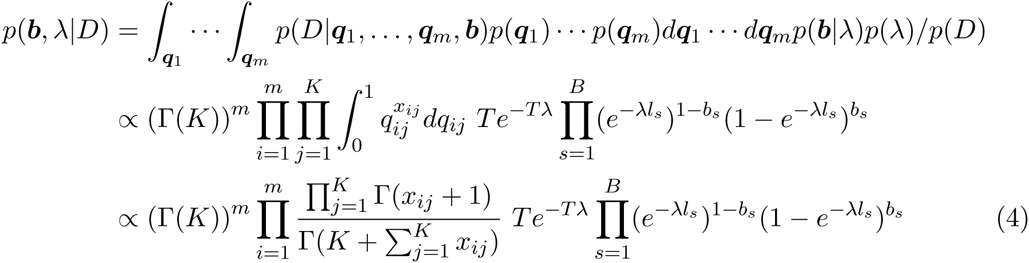

### Inference

We use a MCMC [21] to sample from the posterior distribution of ***b*** and λ. We use a symmetric proposal for ***b*** where the proposed value ***b\**** is the same as ***b*** except for one randomly chosen branch *i* for which 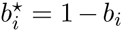. Therefore if the randomly chosen branch *i* holds a changepoint in ***b***, it does not hold a changepoint in ***b\**** and vice versa. To update λ we propose from a normal density with mean equal to the current value of λ and variance equal to 0.1, i.e. λ*|λ ~ *N*(λ,0.1). When the proposed λ* is lower than zero, the move is rejected and the chain stays at λ. The calculation of the Metropolis-Hastings acceptance ratios are given in the supplementary material.

### Model selection

We want to assess whether there is any evidence for differential distribution of phenotype on different parts of the tree. We compare our model (indexed 1) against the null model (indexed 0) of no changepoints on the tree, which is equivalent to λ = 0, by calculating the Bayes factor [19] for the two models. To do this we use reversible jump moves [22] to sample from the joint distribution *p((j,θ*_*j*_)|*D)* where *j* is the index of the model and *θ*_*j*_ is the parameters of model *j.* For a move from null to alternative (0 to 1) model, to match dimensions we generate two random variables *u* and ***v*** and map them such that (λ*, ***b***\*) = (*u*, ***v***). In addition we set the proposal distribution for *u* and ***v***, *q(u*, ***v***) in model 0 to be the same as the prior distribution on λ and ***b*** in model 1. Thus for a proposed move from model 0 to 1 we have:

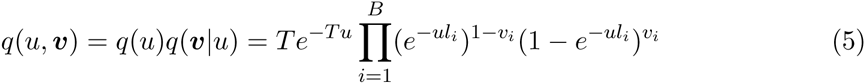

The probability of acceptance of this move is given by:

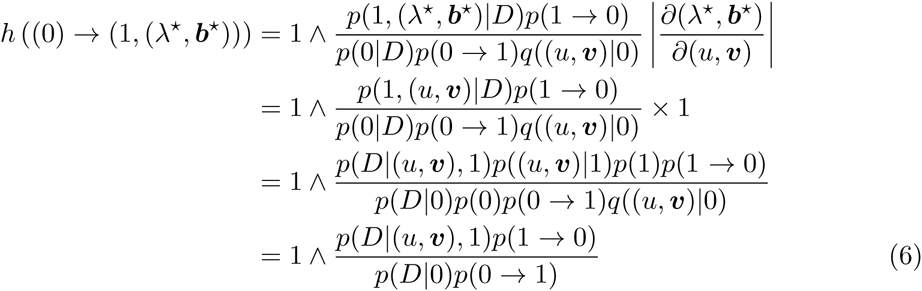

A move from model 1 with parameters (λ, ***b***) to model 0 is made deterministically and is accepted with probability:

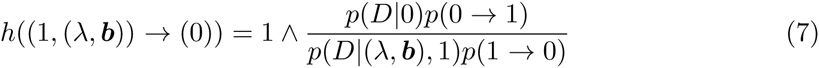

We set *p*(1 → 0) = 0.05 and *p*(0 → 1) = 0.5 and we assume the prior probabilities of the two models are equal p(0) = *p(1) =* 0.5.

### Simulation studies

To investigate the performance of our method, we performed two simulation studies each of which involved repetition over many simulated datasets. In all of these simulations for simplicity we used a binary phenotype and sampled from the posterior distribution of the model parameters using 10^7^ iterations of our MCMC algorithm. All of these simulations were implemented for a single genealogy simulated using the coalescent model [23] with 1000 leaves shown in Figure S1. First we tested how the number of individuals affected by a changepoint and the magnitude of the change in phenotype distribution affects the statistical power to detect a changepoint. Secondly we tested the model selection procedure and the relationship between the posterior expectation of number of changepoints against the true numbers of changepoints. Thirdly we quantified the effect of threshold on the point estimate of ***b***.

## Results

### Simulation study of statistical power

This simulation study was designed to assess the power of the method to detect changepoints on the branches of the tree. The power depends on two factors: the magnitude of the change in the distribution over the phenotype categories which we refer to as *p* and the number of individuals affected by the changepoint which we refer to as *n*. The probability of each phenotype is 0.5 before the changepoint, and after the changepoint the probability of one phenotype increases by *p* whereas the probability of the other phenotype decreases by p. Changepoints with small *p* are difficult to detect as they result in small changes to the observed pattern of distribution of phenotype that are likely to happen by chance alone. Changepoints with small *n* are also difficult to distinguish as lack of data makes the inference more uncertain. We expect that changepoints with large *p* and large *n* to be easier to detect.

The space of *n* × *p* was divided into a grid where *n* = (10, 30, 60, 130,330,500) and *p* = (0.1, 0.2, 0.3, 0.4, 0.5). For each node of the grid (*p*_*i*_, *n*_*j*_) an appropriate branch of the tree shown in Figure S1 was chosen to hold a changepoint, with the remaining branches being left free of changepoints. For each node of the grid we simulated 50 datasets each with a single changepoint. Figure 2 shows for each node of the grid the mean marginal posterior probability of having a changepoint for the branch with the changepoint. A changepoint that causes large changes to the distribution of the phenotype categories and affects a large number of individuals is inferred with a high posterior probability. Changepoints that cause small changes in the distribution or affect few individuals or both result in small posterior probability of having a changepoint.

**Figure 2.**
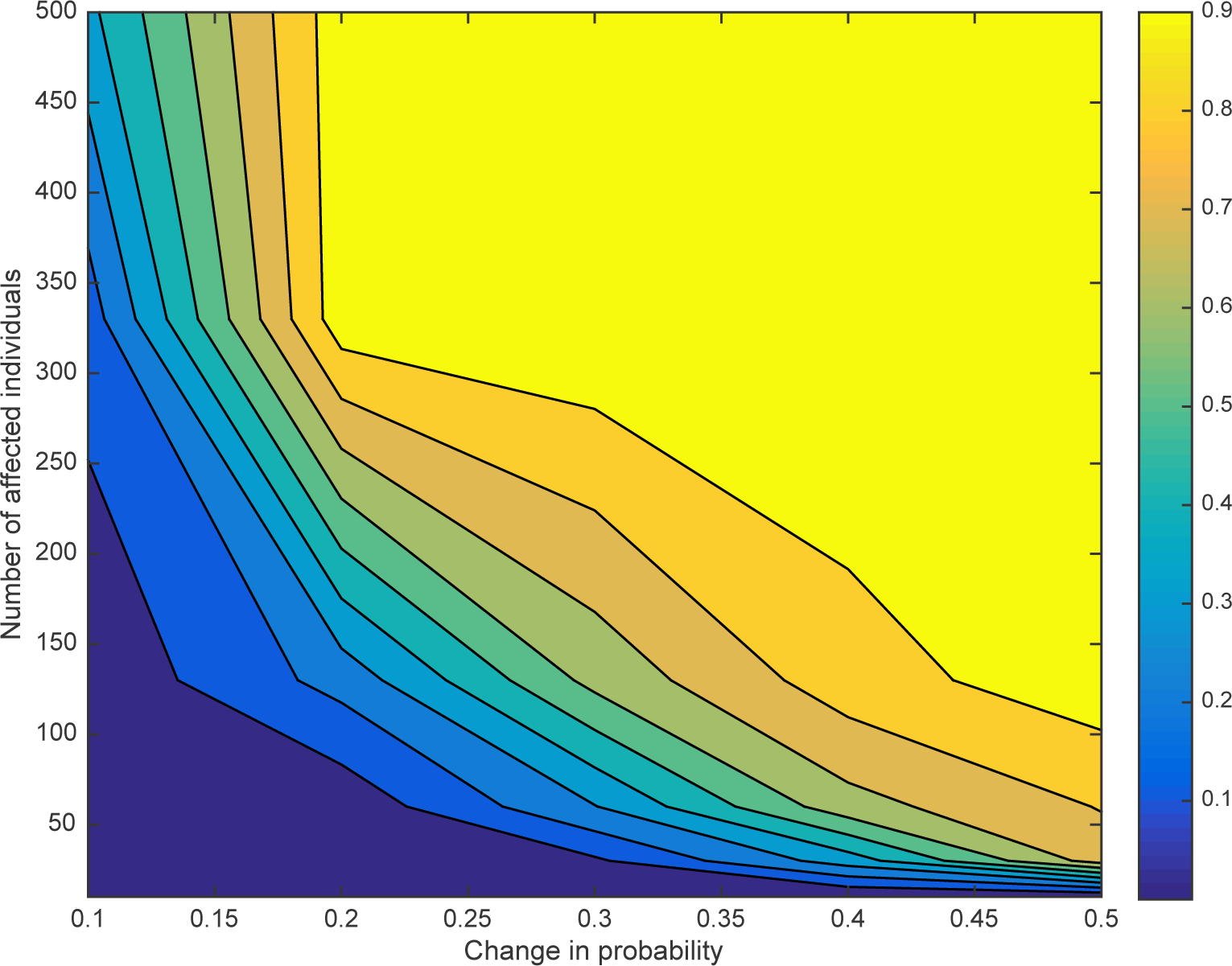
Effect of number of strains and change in distribution. Contour plot of the mean posterior probability of having a changepoint as a function of number *n* of affected individuals and the magnitude *p* of the change in distribution. The space of *n* × *p* was divided into a grid where *n* = (10, 30, 60, 130,330,500) and *p* = (0.1, 0.2, 0.3, 0.4, 0.5).

### Simulation study of model and parameter inference

This simulation study was designed to assess our model selection procedure, the effect of number of changepoints on the inference and the effect of cutoff threshold on the point estimate of ***b***. We simulated 100 datasets for each case of 0, 1, …, 10 changing branches in the tree. The distribution over the phenotypes was uniformly sampled in each case. For each simulated dataset the Bayes factor of our model against null model was estimated (Figure 3A). For the 100 datasets with no changepoint on the tree, all the estimated Bayes factors indicated no significant evidence against the null model (no changepoint on the tree) for any of datasets. Changepoints that result in small changes in the distribution or affect small number of individuals will not be detected. Therefore for some of datasets with a single changing branch there is no significant evidence against the null model, but for some there is strong evidence against the null model. As the number of changing branches on the tree increases, the number of datasets with significant evidence for the alternative model increases. Overall, our method is conservative and should not result in significant evidence for the existence of changepoints unless there is substantial data to support it.

**Figure 3.**
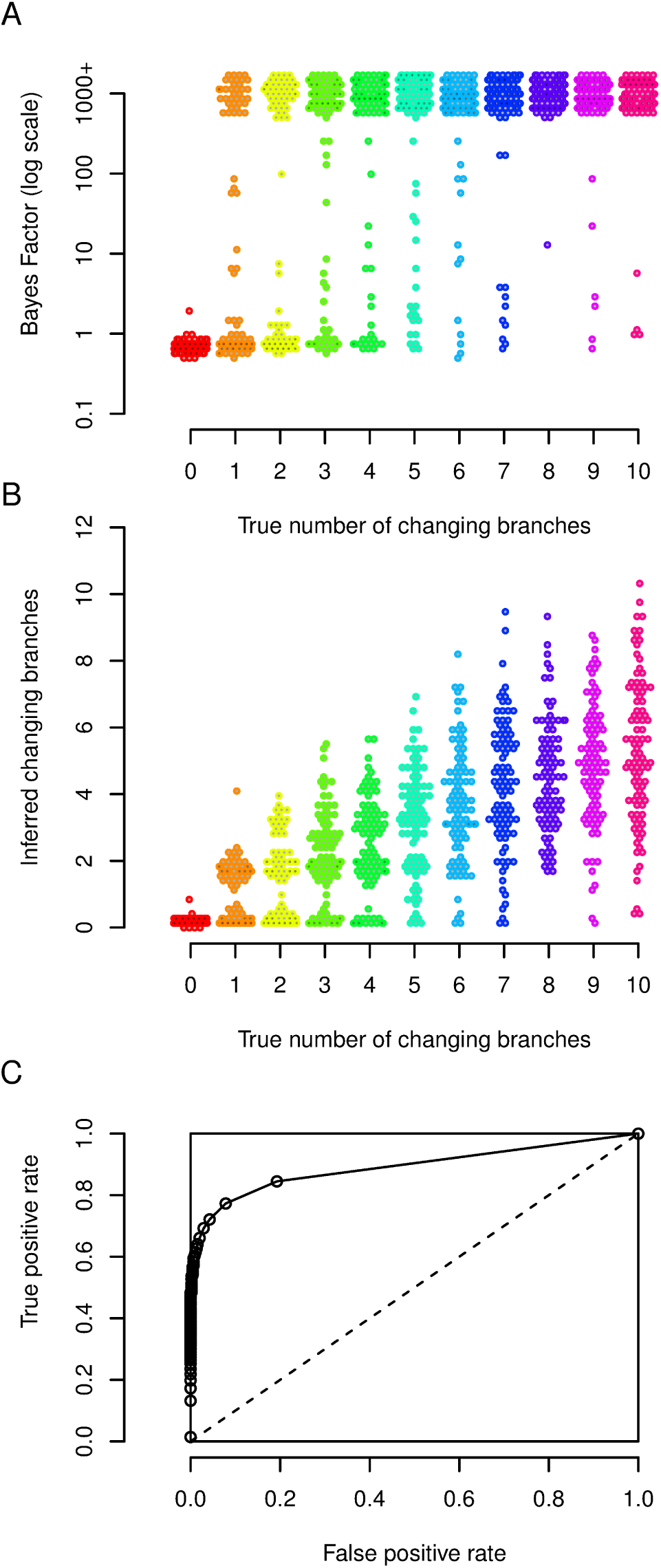
Simulation study of model and parameter inference. (A) Bayes factor values for the changepoint model versus the null model, as a function of the number of changing branches used in the simulation. (B) Distribution of posterior mean number of changing branches as a function of the true number of simulated changing branches. (C) ROC curve: true positive rate as a function of the false positive rate.

Next, we used the simulations to gauge the relationship between the true number of simulated changing branches and its posterior expectation, estimated using Bayesian model averaging [24]. Figure 3B illustrates the results. In the absence of any changepoint, the mean of posterior expectation of number of changing branches is always close to zero. When there are changing branches on the tree, the posterior expectation is downward biased compared to the real value. This is expected as our method cannot detect a changepoint that results in small changes in the distribution or affects few individuals or both. As a result our method is conservative in estimating the number of changepoints on the tree.

In addition we used the simulation results to assess the effect of a cutoff threshold on the point estimate of ***b***. For each of the datasets we inferred a point estimate for ***b*** by applying a threshold to the consensus representation of ***b*** (marginal posterior probability of having a changepoint for each branch of the tree). The threshold was changed from 0 to 1 with increments of 0.01. For each threshold value, the false positive rate and the true positive rate across all of our 1100 simulations was calculated. Figure 3C shows the true positive rate as a function of the false positive rate. This so-called ROC curve has a high area under the curve of 0.891, indicative of good performance of the algorithm [25]. The choice of the cutoff threshold is a trade off between minimising the number of incorrectly inferred changepoints and maximising the number of correctly inferred changepoints. This choice depends on the application and the weight given to sensitivity and specificity in the application.

### Detecting HIV escape mutations from cytotoxic T-lymphocytes

Human leukocyte antigen (HLA) type I genes encode proteins that are present on the surface of almost all human cells. When a cell is infected with a virus, the viral protein is cleaved and small segments of it called epitopes are presented on the cell surface by the HLA encoded proteins. These proteins have a certain amount of affinity and thus in people with the same HLA allele, the same epitope will be recognised and presented on the cell surface. Cytotoxic T lymphocytes (CTLs) are part of the adaptive immune response and recognise these epitopes before destroying the infected cell. A mutation in one of these epitopes can result in no or weak binding of the peptide to the HLA encoded protein or result in lack of recognition by the T cell receptor. Such mutations lead to the virus escaping the immune response of the host. As these mutations can have a fitness cost on transmission to a host with different HLA repertoire, they may revert back to the wild type [26]. Thus the escape mutations on the virus genome are correlated with the host’s HLA alleles.

However to detect these associations one has to account for the possible geographical structuring that could be present in the data. For instance different HCV genotypes are endemic in different parts of the world and HLA allele profiles are also distinct in different populations across the world. When sampling is across different countries or ethnic groups, it is possible that HLA alleles will be associated with specific clusters of the virus simply because of geographical structuring. Several methods have been suggested to account for the non random distribution of HLA alleles on the tips of the phylogenetic tree [27, 28, 29]. We propose that using our algorithm, one can determine if host HLA alleles are randomly distributed on the tips of the virus phylogenetic tree or whether there are clades where the distributions are distinct from each other. The result can then be used to perform stratified association studies conditioned on the clades with distinct HLA distribution.

We used previously published data [30] on a cohort of 261 South Africans to detect HLA-driven evolution of HIV. In this study whole genome viral sequences were aligned and then divided into ten fragments of 1000 nucleotides overlapping by 50 nucleotides. Each partition was then used to produce a maximum likelihood phylogenetic tree. The HLA alleles of the patients were also typed. We used the ten phylogenetic trees from this dataset and the HLA information of the patients as the inputs of our algorithm, considering the presence and absence of each HLA type separately. This resulted in 1197 runs of our software. Figure S2 shows the histogram of the Bayes factors estimated by each run. Only the HLA allele B57 and the tree of the first region of the HIV genome had a Bayes factor conclusively rejecting the null model of no association. Figure 4 shows the distribution of the B57 HLA allele on the tips of the first virus phylogeny. There is a clade of twelve viral individuals where ten of the hosts have the B57 allele, whereas across the rest of the tree there are only seven other hosts with B57 HLA alleles. This clear non random distribution of the HLA B57 could be due to transmission of the virus between closely related people. However we do not detect the same association between the other nine trees from the rest of the genome and HLA B57. An alternative explanation may be that HLA B57 has a significant effect on the evolution of the first 1000 nucleotide of the virus, since HLA B57 is associated with slow progression to disease following HIV infection [31, 32].

**Figure 4.**
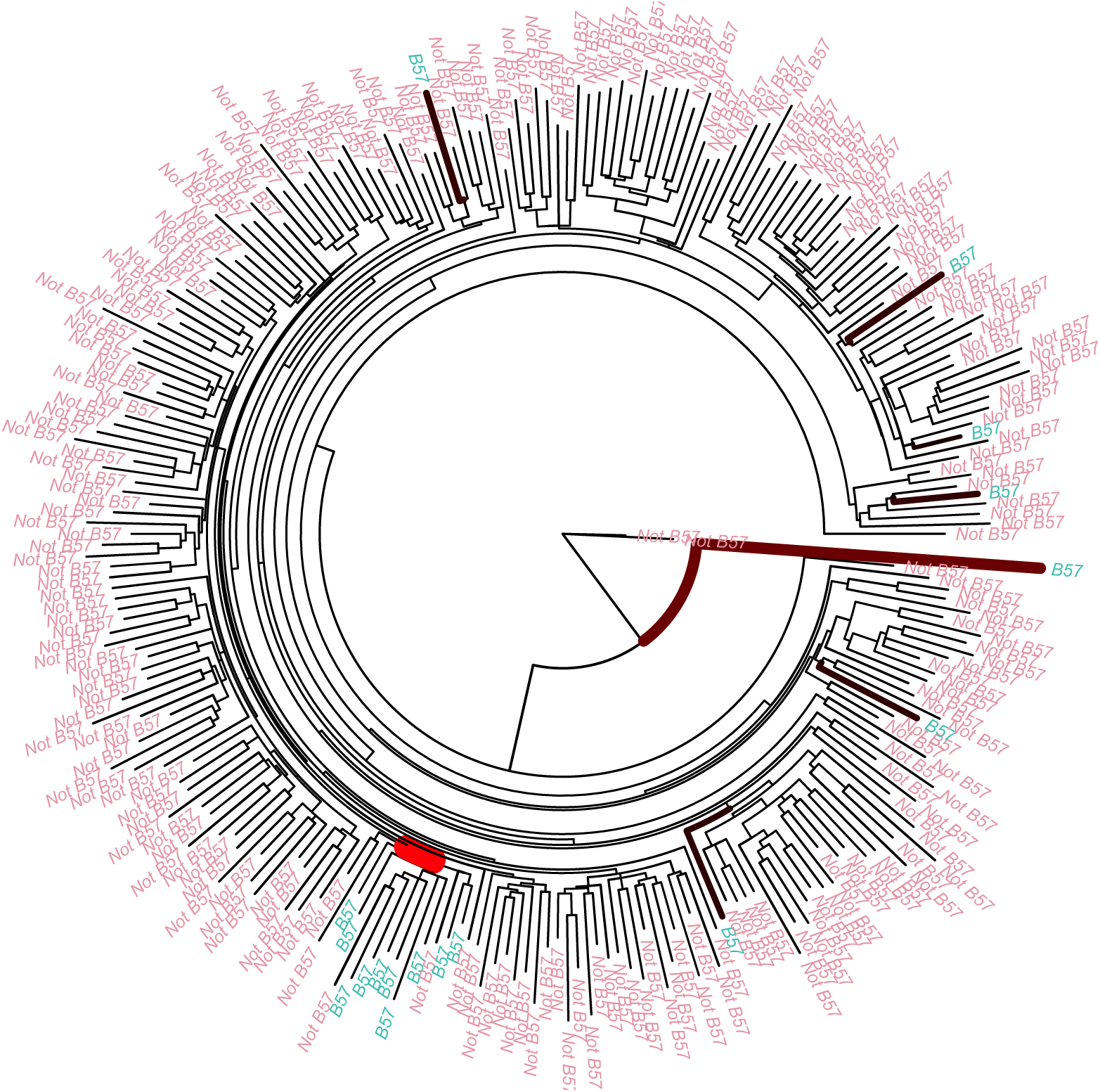
Application to HIV immunology. Phylogenetic tree of 261 HIV infected individuals from the first 1000 nucleotides with the tips coloured according to presence and absence of HLA B57 in the host. The thickness and colour of the branches are proportional to the posterior probability of having a changepoint.

### Inferring host range within a lineage of Salmonella enterica

*Salmonella enterica* is a bacterial pathogen made of multiple lineages with different host ranges [33, 34, 35]. Many lineages can infect a wide range of animals, whereas some are mostly found in specific hosts and yet others have become restricted to a single host type, for example the Typhi and Paratyphi A lineages which evolved in convergence towards infecting only humans [36]. The Typhimurium DT104 lineage has been responsible for a global multidrug resistant epidemic since the 1990s in both humans and farm animals [37, 38, 39]. Typhimurium DT104 can infect both animals and humans, but it is unclear if there are sublineages within DT104 that infect one host type more than the other, and to what extent the epidemics in animals and humans are associated. Traditional molecular typing techniques do not provide enough genetic resolution to answer this question. A recent study sequenced the whole genomes of 142 human strains and 120 animal strains isolated in Scotland between 1990 and 2011 [40]. A maximum-likelihood tree was computed based on the non-recombinant core genome using RAxML [41] and here we applied our algorithm to this tree, using animal versus human source as the phenotype.

The null model of random distribution of hosts around the tree was decisively rejected in favour of the changepoint model, with the reversible jump MCMC never exploring the null model after initial burnin. The posterior mean number of changing branches was 9.7, with 95% credibility interval ranging from 5 to 16. Changes in the host range were especially evident on four branches (Figure 5), corresponding to posterior probabilities of 99%, 95%, 90% and 72%, with two further branches with probability 54%, one with 39% and all others below 20%. Amongst the four branches with highest support, the oldest corresponds to an increase in the frequency of infection of animals for a large clade of 265 isolates within DT104. The other three branches all occurred within this clade, and correspond to three separate further increase in the frequency of infection of animals for three subclades containing 12, 15 and 59 isolates, respectively. These results confirm and refine the original conclusions of the study in which the data was presented [40], that the epidemic of DT104 in Scotland was not homogenous in humans and animals. Specifically, a sublineage increasingly became restricted to infecting only animals and not humans, which could be the result of either adaptation or niche segregation.

**Figure 5.**
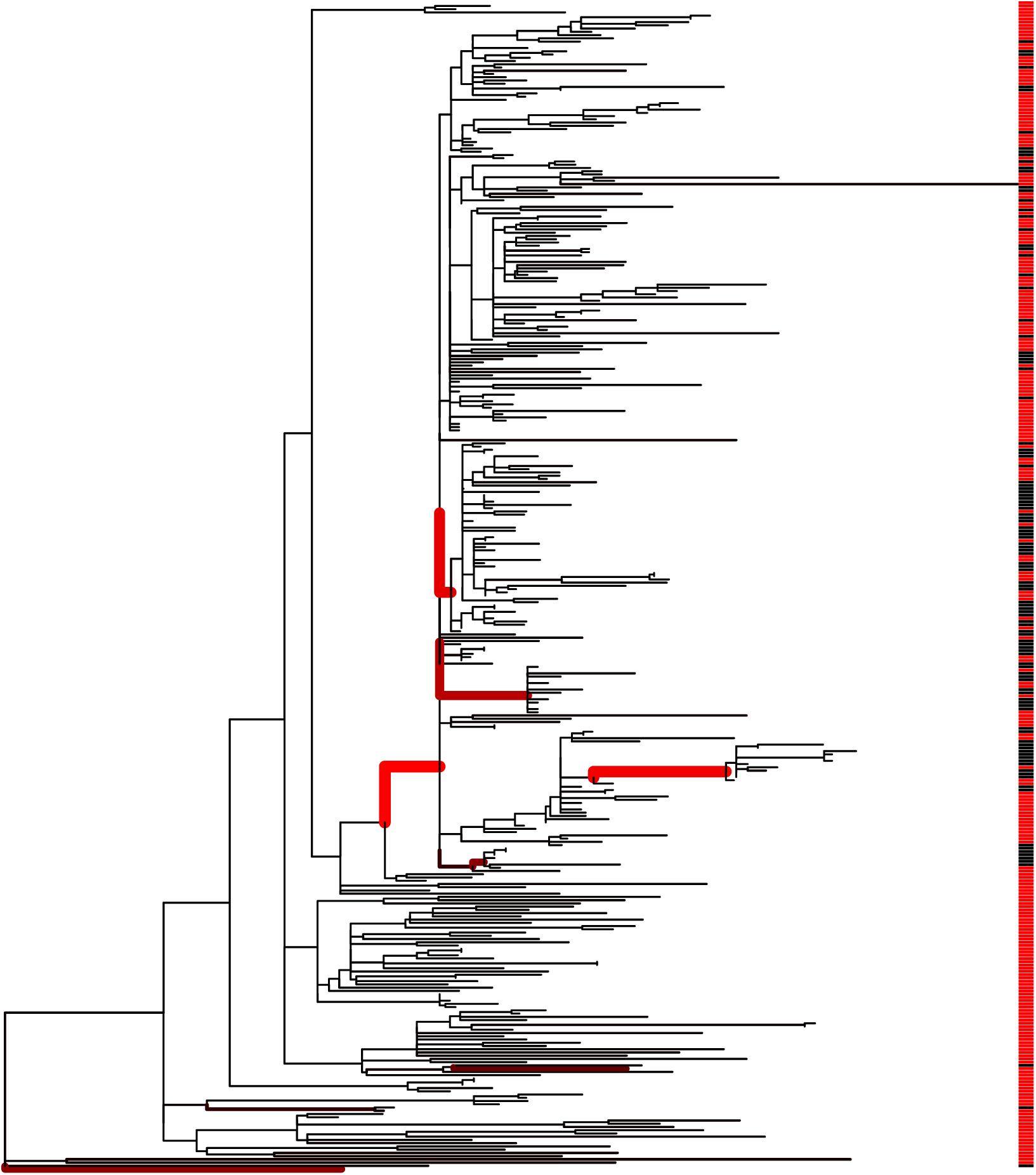
Application to Salmonella ecology. Maximum-likelihood phylogenetic tree from a previous study of Typhimurium DT104 [40], with the color on the right indicating the isolates came from either human (red) or animal (black) sources. The results of our algorithm are shown by the thickness and redness of the branches, which are both proportional to the posterior probability of host range change on the given branch.

## Discussion

This study is based on the concept of phenotype distribution, which is the distribution of phenotypes that a given genotype may express depending on environmental factors, as a result of phenotypic plasticity [2, 3]. We presented a model in which the phenotype distribution is allowed to change along the branches of a phylogenetic tree, and an efficient Bayesian method to perform inference under this model. Given phenotype observations for the leaves of a phylogeny, we showed that our method can be used to detect branches on which the phenotype distribution changed significantly. Consequently, a phylogeny can be demarcated into lineages with distinct phenotype distributions.

There are many ways in which our approach could be extended, for example to be applicable to continuous rather than categorical phenotype measurements, or to allow the evolution of the phenotype distribution to be more progressive, for example by making this distribution after a changepoint correlated with, rather than independent from, the distribution before the changepoint. We did not attempt to model the potential for error in either the input phylogeny or input phenotype measurements. Uncertainty about the tree could be accounted for by applying our method to a sample of trees from the posterior distribution of the trees that are produced by Bayesian phylogenetic software such as MrBayes and BEAST [42, 43]. However, we expect that a little inaccuracy in the tree would not drastically affect the result of our method, and likewise for the phenotype measurement, because the results depend on phenotype distributions which are themselves stochastic. This is unlike methods that consider changes in the phenotype itself, such as ancestral state reconstructions [15], for which a mistake in a single phenotype measurement implies an additional evolutionary event for the phenotype. When considering phenotypes with imperfect heritability [1], we argue that modelling the evolution of the phenotype distribution is more biologically relevant than modelling the evolution of the phenotype measurement.

There are many research areas in which the method we proposed could be useful, and we presented two examples in HIV immunology and bacterial ecology. For example, our approach could help provide a definition of microbial species. Detecting incipient speciation requires to distinguish between ecologically distinct populations in the same community [44, 45, 46]. In this case the phenotype would be ecological or pathogenicity measurements, and the aim is to determine if different phylogenetic clades have distinct distributions over the measurable ecological quantities [47, 48]. Another potential area of application is genome wide association studies (GWAS) in organisms that reproduce clonally. Population structure is a confounding effect in GWAS [49] and this is especially important for clonal organisms [50]. One way to account for this population structure would be to use our method to find the clades on the phylogenetic tree where the phenotype of interest is uniquely distributed and perform GWAS stratified by those clusters.

## Acknowledgements

This research was made possible by a James Martin fellowship awarded to M. A. Ansari. X. Didelot received funding from the Biotechnology and Biological Sciences Research Council (BB/L023458/1) and the National Institute for Health Research (HPRU-2012-10080). The authors would like to thank Remi Bardenet, Daniel Falush, Philip Maybank and Gil McVean for their insightful discussions.

